# A framework for effective application of machine learning to microbiome-based classification problems

**DOI:** 10.1101/816090

**Authors:** Begüm D. Topçuoğlu, Nicholas A. Lesniak, Mack Ruffin, Jenna Wiens, Patrick D. Schloss

**Affiliations:** Department of Microbiology and Immunology, University of Michigan, Ann Arbor, MI 48109; Department of Electrical Engineering and Computer Science, University of Michigan, Ann Arbor, MI 48109; Department of Family Medicine and Community Medicine, Penn State Hershey Medical Center, Hershey, PA

## Abstract

Machine learning (ML) modeling of the human microbiome has the potential to identify microbial biomarkers and aid in the diagnosis of many diseases such as inflammatory bowel disease, diabetes, and colorectal cancer. Progress has been made towards developing ML models that predict health outcomes using bacterial abundances, but inconsistent adoption of training and evaluation methods call the validity of these models into question. Furthermore, there appears to be a preference by many researchers to favor increased model complexity over interpretability. To overcome these challenges, we trained seven models that used fecal 16S rRNA sequence data to predict the presence of colonic screen relevant neoplasias (SRNs; n=490 patients, 261 controls and 229 cases). We developed a reusable open-source pipeline to train, validate, and interpret ML models. To show the effect of model selection, we assessed the predictive performance, interpretability, and training time of L2-regularized logistic regression, L1 and L2-regularized support vector machines (SVM) with linear and radial basis function kernels, decision trees, random forest, and gradient boosted trees (XGBoost). The random forest model performed best at detecting SRNs with an AUROC of 0.695 [IQR 0.651-0.739] but was slow to train (83.2 h) and not inherently interpretable. Despite its simplicity, L2-regularized logistic regression followed random forest in predictive performance with an AUROC of 0.680 [IQR 0.625-0.735], trained faster (12 min), and was inherently interpretable. Our analysis highlights the importance of choosing an ML approach based on the goal of the study, as the choice will inform expectations of performance and interpretability.

**Importance:** Diagnosing diseases using machine learning (ML) is rapidly being adopted in microbiome studies. However, the estimated performance associated with these models is likely over-optimistic. Moreover, there is a trend towards using black box models without a discussion of the difficulty of interpreting such models when trying to identify microbial biomarkers of disease. This work represents a step towards developing more reproducible ML practices in applying ML to microbiome research. We implement a rigorous pipeline and emphasize the importance of selecting ML models that reflect the goal of the study. These concepts are not particular to the study of human health but can also be applied to environmental microbiology studies.

## Background

As the number of people represented in human microbiome datasets grow, there is an increasing desire to use microbiome data to diagnose diseases. However, the structure of the human microbiome is remarkably variable among individuals to the point where it is often difficult to identify the bacterial populations that are associated with diseases using traditional statistical models. For example it is not possible to classify individuals as having healthy colons or screen relevant neoplasia using Bray-Curtis distances based on the 16S rRNA gene sequences collected from fecal samples [Figure S1]. This variation is likely due to the ability of many bacterial populations to fill the same niche such that different populations cause the same disease in different individuals. Furthermore, a growing number of studies have shown that it is rare for a single bacterial species to be associated with a disease. Instead, subsets of the microbiome account for differences in health. Traditional statistical approaches do not adequately account for the variation in the human microbiome and typically consider the protective or risk effects of each bacterial population separately (1). Recently, machine learning (ML) models have grown in popularity among microbiome researchers as our ability to sample large numbers of individuals has grown; such models can effectively account for the interpersonal microbiome variation and the ecology of disease because they consider the relative abundance of each bacterial population in the context of others rather than in isolation.

ML models can be used to increase our understanding of the variation in the structure of existing data and in making predictions about new data. Researchers have used ML models to diagnose and understand the ecological basis of diseases such as liver cirrhosis, colorectal cancer, inflammatory bowel diseases, obesity, and type 2 diabetes (2–19). The task of diagnosing an individual relies on a rigorously validated model. However, there are common methodological and reporting problems that arise when applying ML to such data that need to be addressed for the field to progress. These problems include a lack of transparency in which methods are used and how these methods are implemented; evaluating models without separate held-out test data; unreported variation between the predictive performance on different folds of cross-validation; and unreported variation between cross-validation and testing performances. Though the microbiome field is making progress to avoid some of these pitfalls including validating their models on independent datasets (8, 19, 20) and introducing accessible and open-source ML tools (21–24), more work is needed to improve reproducibility further and minimize overestimating for model performance.

Among microbiome researchers, the lack of justification when selecting a modeling approach has often been due to an implicit assumption that more complex models are better. This has resulted in a trend towards using non-linear models such as random forest and deep neural networks (3, 12, 25–27) over simpler models such as logistic regression or other linear models (19, 23, 28). Although in some cases, complex models may capture important non-linear relationships and therefore yield better predictions, they can also result in black boxes that lack interpretability. Such models require post hoc explanations to quantify the importance of each feature in making predictions. Depending on the goal of the modeling, other approaches may be more appropriate. For example, researchers trying to identify the microbiota associated with disease may desire a more interpretable model, whereas clinicians may emphasize predictive performance. Nonetheless, it is essential to understand that the benefit of more complex, less interpretable models may be minimal (29–31). It is important for researchers to justify their choice of modeling approach.

In this study, we provided steps toward standardization of machine learning methods for microbiome studies which are often poorly documented and executed. To showcase a rigorous ML pipeline and to shed light on how ML model selection can affect modeling results, we performed an empirical analysis comparing the predictive performance, interpretability, data requirements, and training times of seven modeling approaches with the same dataset and pipeline. We built three linear models with different forms of regularization: L2-regularized logistic regression and L1 and L2-regularized support vector machines (SVM) with a linear kernel. We also trained four non-linear models: SVM with radial basis function kernel, a decision tree, random forest, and gradient boosted trees. We compared their predictive performance, interpretability, and training time. To demonstrate the performance of these modeling approaches and our pipeline, we present a case study using data from a previously published study that sought to classify individuals as having healthy colons or colonic lesions based on the 16S rRNA gene sequences collected from fecal samples (4). This dataset was selected because it is a relatively large collection of individuals (N=490) connected to a clinically significant disease where there is ample evidence that the disease is driven by variation in the microbiome (2, 4, 5, 32). With this dataset, we developed an ML pipeline that can be used in many different scenarios for training and evaluating models. This framework can be easily applied to other host-associated and environmental microbiome datasets. We also provided an aspirational rubric for evaluating the rigor of ML practices applied to microbiome data [Table S1] to urge the audience to be diligent in their study design and model selection, development, evaluation, and interpretation.

## Results

### Model selection and pipeline construction

We established a reusable ML pipeline for model selection and evaluation, focusing on seven different commonly used supervised learning algorithms [Figure 1, Table 1].

**Table 1.**
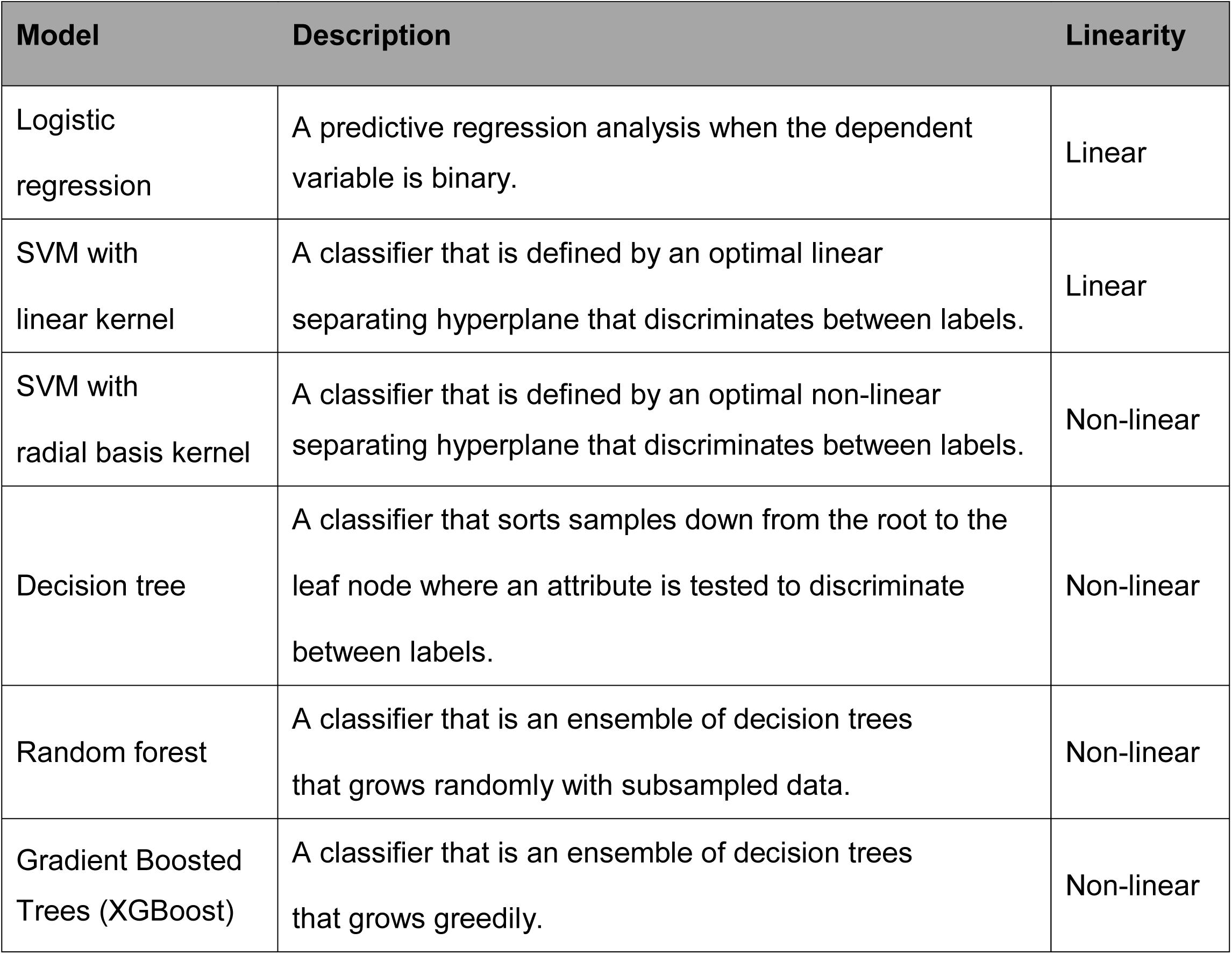
Characteristics of machine learning models in this study.

| Model | Description | Linearity |
| --- | --- | --- |
| Logistic regression | A predictive regression analysis when the dependent variable is binary. | Linear |
| SVM with linear kernel | A classifier that is defined by an optimal linear separating hyperplane that discriminates between labels. | Linear |
| SVM with radial basis kernel | A classifier that is defined by an optimal non-linear separating hyperplane that discriminates between labels. | Non-linear |
| Decision tree | A classifier that sorts samples down from the root to the leaf node where an attribute is tested to discriminate between labels. | Non-linear |
| Random forest | A classifier that is an ensemble of decision trees that grows randomly with subsampled data. | Non-linear |
| Gradient Boosted Trees (XGBoost) | A classifier that is an ensemble of decision trees that grows greedily. | Non-linear |

**Figure 1.**
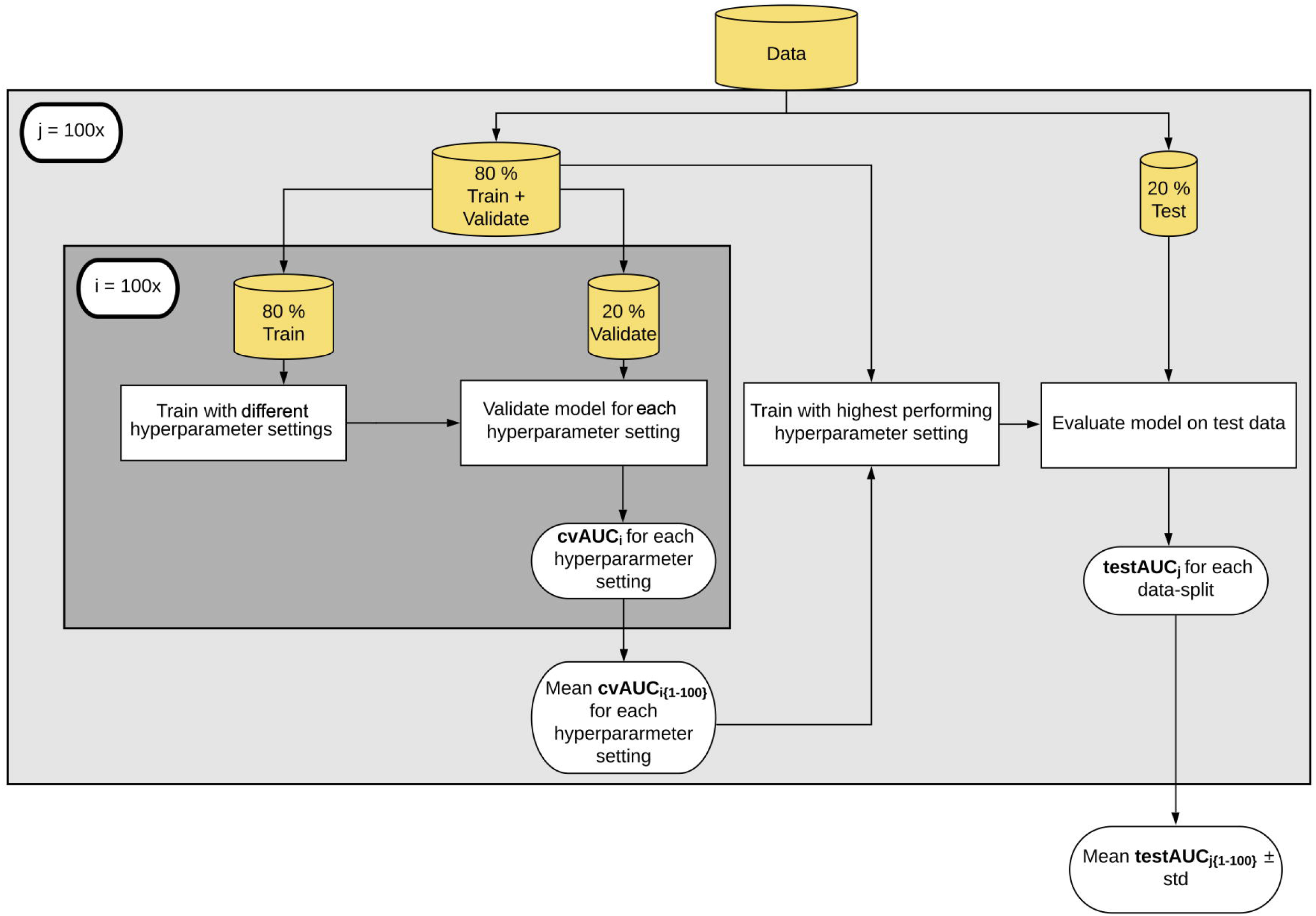
Machine learning pipeline. We split the data to create a training (80%) and held-out test set (20%). The splits were stratified to maintain the overall class distribution. We performed five-fold cross-validation on the training data to select the best hyperparameter setting and then used these hyperparameters to train the models. The model was evaluated on the held-out data set. Abbreviations: cvAUC, cross-validation area under the receiver operating characteristic curve.

First, we randomly split the data into training and test sets so that the training set consisted of 80% of the full dataset, while the test set was composed of the remaining 20% [Figure 1]. To maintain the distribution of controls and cases found in the full dataset, we performed stratified splits. For example, our full dataset included 490 individuals. Of these, 261 had healthy colons (53%) and 229 had a screen relevant neoplasia (SRN; 46.7%). A training set included 393 individuals, of which 209 had an SRN (53%), while the test set was composed of 97 individuals, of which 52 had an SRN (54%). The training data were used to build and select the models, and the test set was used for evaluating the model. We trained seven different models using the training data [Table 1].

Model selection requires tuning hyperparameters. Hyperparameters are parameters that need to be specified or tuned by the user, in order to train a model for a specific modeling problem. For example, when using regularization, C is a hyperparameter that indicates the penalty for overfitting. Hyperparameters are tuned using the training data to find the best model. We selected hyperparameters by performing repeated five-fold cross-validation (CV) on the training set [Figure 1]. The five-fold CV was also stratified to maintain the overall case and control distribution. We chose the hyperparameter values that led to the best average CV predictive performance using the area under the receiver operating characteristic curve (AUROC) [Figure S2 and S3]. The AUROC ranges from 0, where the model’s predictions are perfectly incorrect, to 1.0, where the model perfectly distinguishes between cases and controls. An AUROC value of 0.5 indicates that the model’s predictions are no different than random. To select hyperparameters, we performed a grid search for hyperparameter settings when training the models. Default hyperparameter settings in developed ML packages available in R, Python, and MATLAB programming languages may be inadequate for effective application of classification algorithms and need to be optimized for each new ML task. For example, L1-regularized SVM with linear kernel showed large variability between different regularization strengths (C) and benefited from tuning as the default C parameter was 1 [Figure S2].

Once hyperparameters were selected, we trained the model using the full training dataset and applied the final model to the held-out data to evaluate the testing predictive performance of each model. The data-split, hyperparameter selection, training and testing steps were repeated 100 times to obtain a robust interpretation of model performance, less likely to be affected by a “lucky” or “unlucky” split [Figure 1].

### Predictive performance and generalizability of the seven models

We evaluated the predictive performance of the seven models to classify individuals as having healthy colons or SRNs [Figure 2]. The predictive performance of random forest model was higher than other ML models with a median 0.695 [IQR 0.650-0.739], though not significantly (p=0.5; The p-value was manually calculated using the sampling distribution of the test statistic under the null hypothesis) (Figure S4). Similarly, L2-regularized logistic regression, XGBoost, L2-regularized SVM with linear and radial basis function kernel AUROC values were not significantly different from one another and had median AUROC values of 0.680 [IQR 0.639-0.750], 0.679 [IQR 0.643-0.746], 0.678 [IQR 0.639-0.750] and 0.668 [IQR 0.639-0.750], respectively. L1-regularized SVM with linear kernel and decision tree had significantly lower AUROC values than the other ML models with median AUROC of 0.650 [IQR 0.629-0.760] and 0.601 [IQR 0.636-0.753], respectively [Figure 2]. Interestingly, these results demonstrate that the most complex model (XGBoost) did not have the best performance and that the most interpretable models (L2-regularized logistic regression and L2-regularized SVM with linear kernel) performed nearly as well as non-linear models.

**Figure 2.**
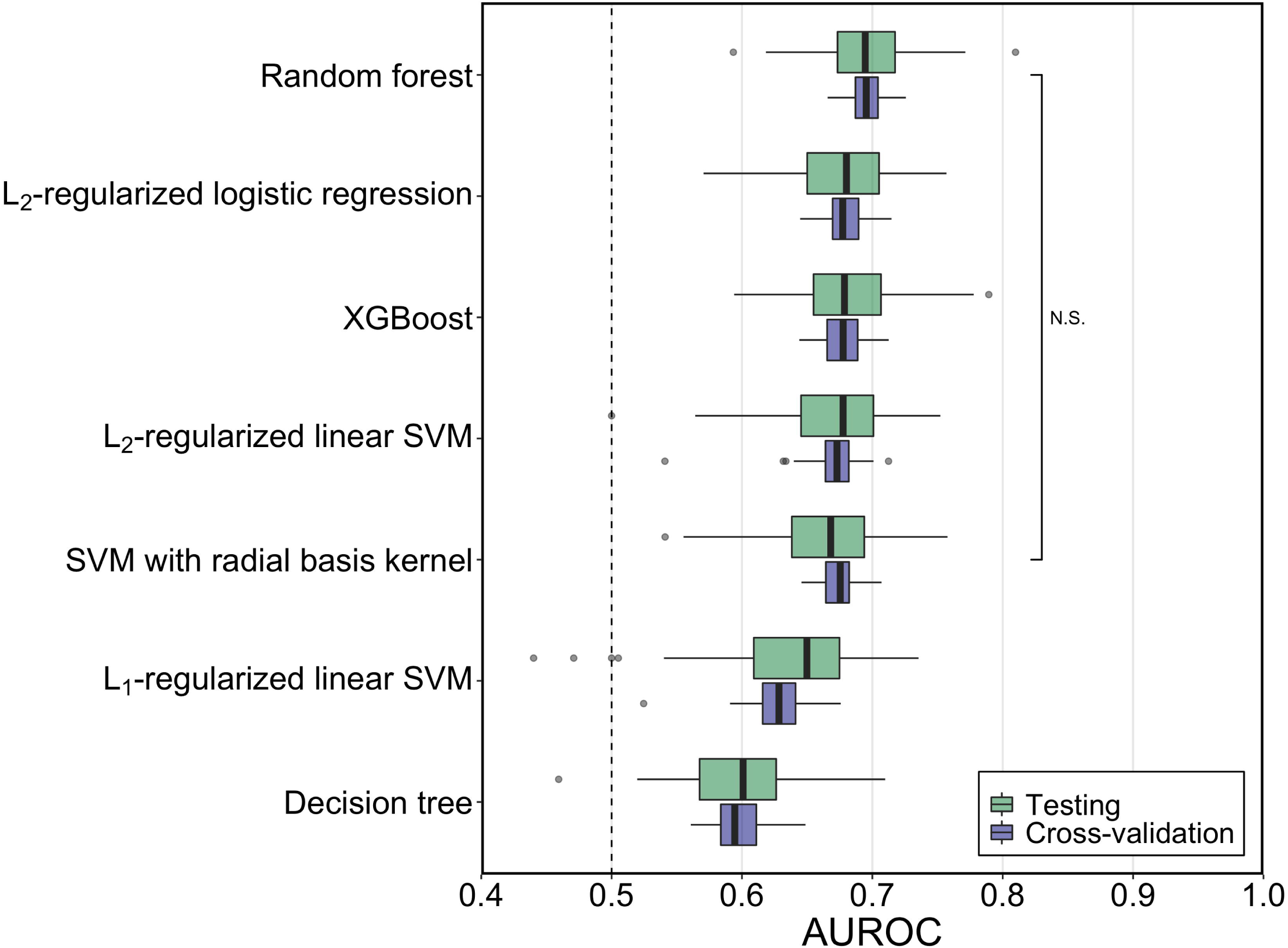
Generalization and classification performance of ML models using AUROC values of all cross-validation and testing performances. The median AUROC for diagnosing individuals with SRN using bacterial abundances was higher than chance (depicted by a horizontal line at 0.50) for all the ML models. The predictive performance of random forest model was higher than other ML models, though not significantly (p > 0.05). L2-regularized logistic regression, XGBoost, L2-regularized SVM with linear and radial basis function kernel performances were not significantly different from one another. The boxplot shows quartiles at the box ends and the median as the horizontal line in the box. The whiskers show the farthest points that were not outliers. Outliers were defined as those data points that are not within 1.5 times the interquartile ranges.

To evaluate the generalizability of each model, we compared the median cross-validation AUROC to the median testing AUROC. If the difference between the cross-validation and testing AUROCs was large, then that could indicate that the models were overfit to the training data. The largest difference in median AUROCs was 0.021 in L1-regularized SVM with linear kernel, followed by SVM with radial basis function kernel and decision tree with a difference of 0.007 and 0.006, respectively [Figure 2]. These differences were relatively small and gave us confidence in our estimate of the generalization performance of the models.

To evaluate the variation in the estimated performance, we calculated the range of AUROC values for each model using 100 data-splits. The range among the testing AUROC values within each model varied by 0.230 on average across the seven models. If we had only done a single split, then there is a risk that we could have gotten lucky or unlucky in estimating model performance. For instance, the lowest AUROC value of the random forest model was 0.593, whereas the highest was 0.810. These results showed that depending on the data-split, the testing performance can vary [Figure 2]. Therefore, it is important to employ multiple data splits when estimating generalization performance.

To show the effect of sample size on model generalizability, we compared cross-validation AUROC values of L2-regularized logistic regression and random forest models when we subsetted our original study design with 490 subjects to 15, 30, 60, 120, and 245 subjects [Figure S5]. The variation in cross-validation performance within both models at smaller sample sizes was larger than when the full collection of samples was used to train and validate the models. Because of the high dimensionality of the microbiome data (6920 OTUs), large sample sizes can lead to better models.

### Interpretation of each ML model

We often use ML models not just to predict a health outcome, but also to identify potential biomarkers for disease. Therefore, model interpretation becomes crucial for microbiome studies. Interpretability is related to the degree to which humans can understand the reasons behind a model prediction (33–35). ML models often decrease in interpretability as they increase in complexity. In this study, we used two methods to help interpret our models.

First, we interpreted the feature importance of the linear models (L1 and L2-regularized SVM with linear kernel and L2-regularized logistic regression) using the median rank of absolute feature weights for each OTU [Figure 3]. We also reviewed the signs of feature weights to determine whether an OTU was associated with classifying a subject as being healthy or having an SRN. It was encouraging that many of the highest-ranked OTUs were shared across these three models (e.g., OTUs 50, 426, 609, 822, 1239). The benefit of this approach was knowing the sign and magnitude of each OTU coefficient in the trained model. This allowed us to immidiately interpret negative and positive coefficient signs as protective and risk factors, respectively and the magnitude as the impact of these factors. However, this approach is limited to linear models or models with prespecified interaction terms.

**Figure 3.**
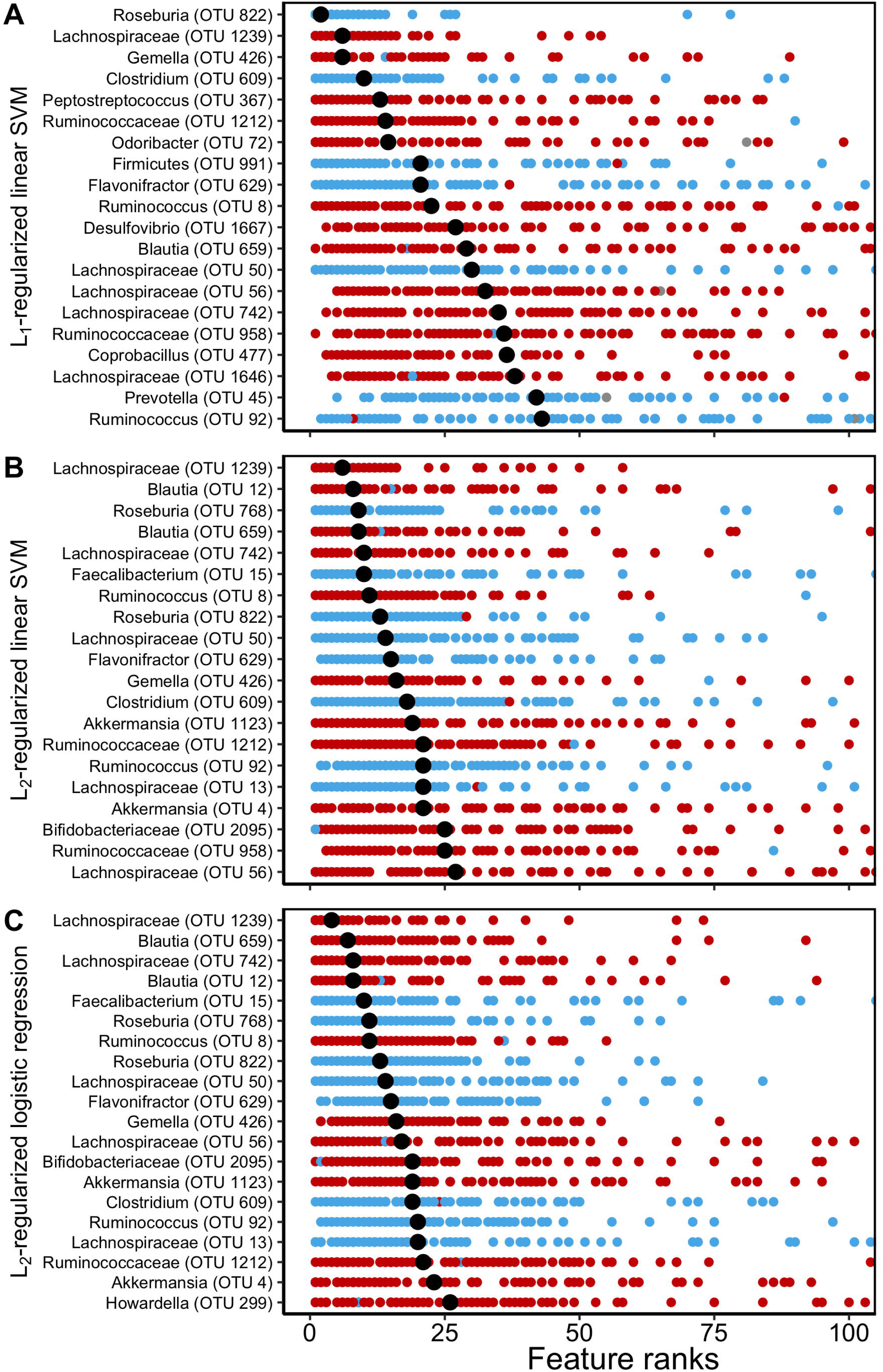
Interpretation of the linear ML models. The ranks of absolute feature weights of (A) L1-regularized SVM with linear kernel, (B) L2-regularized SVM with linear kernel, and (C) L2- regularized logistic regression, were ranked from highest rank, 1, to lowest rank, 100, for each data-split. The feature ranks of the 20 highest ranked OTUs based on their median ranks (median shown in black) are reported here. OTUs that were associated with classifying a subject as being healthy had negative signs and were shown in blue. OTUs that were associated with classifying a subject having an SRN had positive signs and were shown in red.

Second, to analyze non-linear models, we interpreted the feature importance using permutation importance (36). Whereas the absolute feature weights were determined from the trained models, here we measured importance using the held-out test data. Permutation importance analysis is a post hoc explanation of the model, in which we randomly permuted groups of perfectly correlated features together and other features individually across the two groups in the held-out test data [Figure S6]. We then calculated how much the predictive performance of the model (i.e., testing AUROC values) decreased when each OTU or group of OTUs was randomly permuted. We ranked the OTUs based on how much the median testing AUROC decreased when it was permuted; the OTU with the largest decrease ranked highest [Figure 4]. Among the twenty OTUs with the largest impact, there was only one OTU (OTU 822) that was shared among all of the models; however, we found three OTUs (OTUs 58, 110, 367) that were important in each of the tree-based models. Similarly, the random forest and XGBoost models shared four of the most important OTUs (OTUs 2, 12, 361, 477). Permutation analysis results also revealed that with the exception of the decision tree model, removal of any individual OTU had minimal impact on model performance. For example, if OTU 367 was permuted across the samples in the decision tree model, the median AUROC dropped from 0.601 to 0.525. In contrast, if the same OTU was permuted in the random forest model, the AUROC only dropped from 0.695 to 0.680, which indicated high degree of collinearity in the dataset. Permutation analysis allowed us to gauge the importance of each OTU in non-linear models and partially account for collinearity by grouping correlated OTUs to determine their impact as a group.

**Figure 4.**
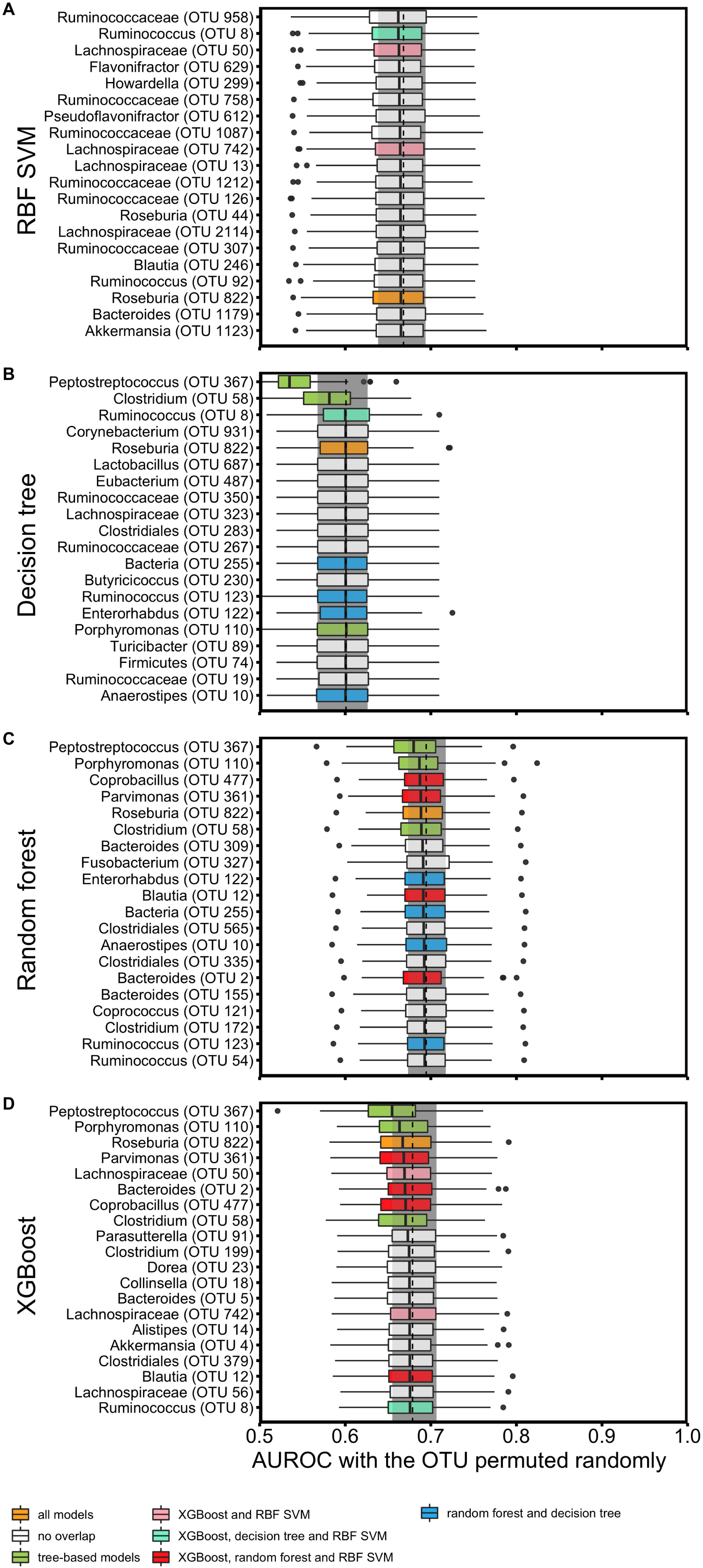
Interpretation of the non-linear ML models. (A) SVM with radial basis kernel, (B) decision tree, (C) random forest, and (D) XGBoost feature importances were explained using permutation importance on the held-out test data set. The gray rectangle and the dashed line show the IQR range and median of the base testing AUROC without any permutation. The 20 OTUs that caused the largest decrease in the AUROC when permuted are reported here. The colors of the box plots represent the OTUs that were shared among the different models; yellow were OTUs that were shared among all the non-linear models, green were OTUs that were shared among the tree-based models, green were the OTUs shared among SVM with radial basis kernel, decision tree and XGBoost, pink were the OTUs shared among SVM with radial basis kernel and XGBoost only, red were the OTUs shared among random forest and XGBoost only and blue were the OTUs shared among decision tree and random forest only. For all of the tree-based models, a *Peptostreptococcus* species (OTU00367) had the largest impact on predictive performance.

To further highlight the differences between the two interpretation methods, we used permutation importance to interpret the linear models [Figure S7]. When we analyzed the L1- regularized SVM with linear kernel model using feature rankings based on weights [Figure 3] and permutation importance [Figure S7], 17 of the 20 top OTUs (e.g., OTU 609, 822, 1239) were deemed important by both interpretation methods. Similarly, for the L2-regularized SVM and L2-regularized logistic regression, 9 and 12 OTUs, respectively, were shared among the two interpretation methods. These results indicate that both methods are consistent in selecting the most important OTUs.

We also compared the top 20 OTUs selected by permutation importance in L2-regularized logistic regression [Figure S7] and the highest performing tree-based models, random forest and XGBoost [Figure 4]. Two and five OTUs, respectively, were shared among the models. These results indicate that we were able to identify important OTUs that are shared across the highest performing linear and non-linear models when we use permutation importance as our interpretation method.

We then evaluated the difference in relative abundances of the top 20 OTUs identified in L2- regularized logistic regression and random forest models between healthy patients and patients with SRNs [Figure S8]. There were minimal differences in the median relative abundances across OTUs between different diagnoses. This supports our claim that it is not possible to differentiate disease versus healthy states by focusing on individual taxa. The ability for ML models to simultaneously consider the relative abundances of multiple OTUs and their context dependency is a great advantage over traditional statistical approaches that consider each OTU in isolation.

### The computational efficiency of each ML model

We compared the training times of the seven ML models. The training times increased with the complexity of the model and the number of potential hyperparameter combinations. Also, the linear models trained faster than non-linear models [Figure 5].

**Figure 5.**
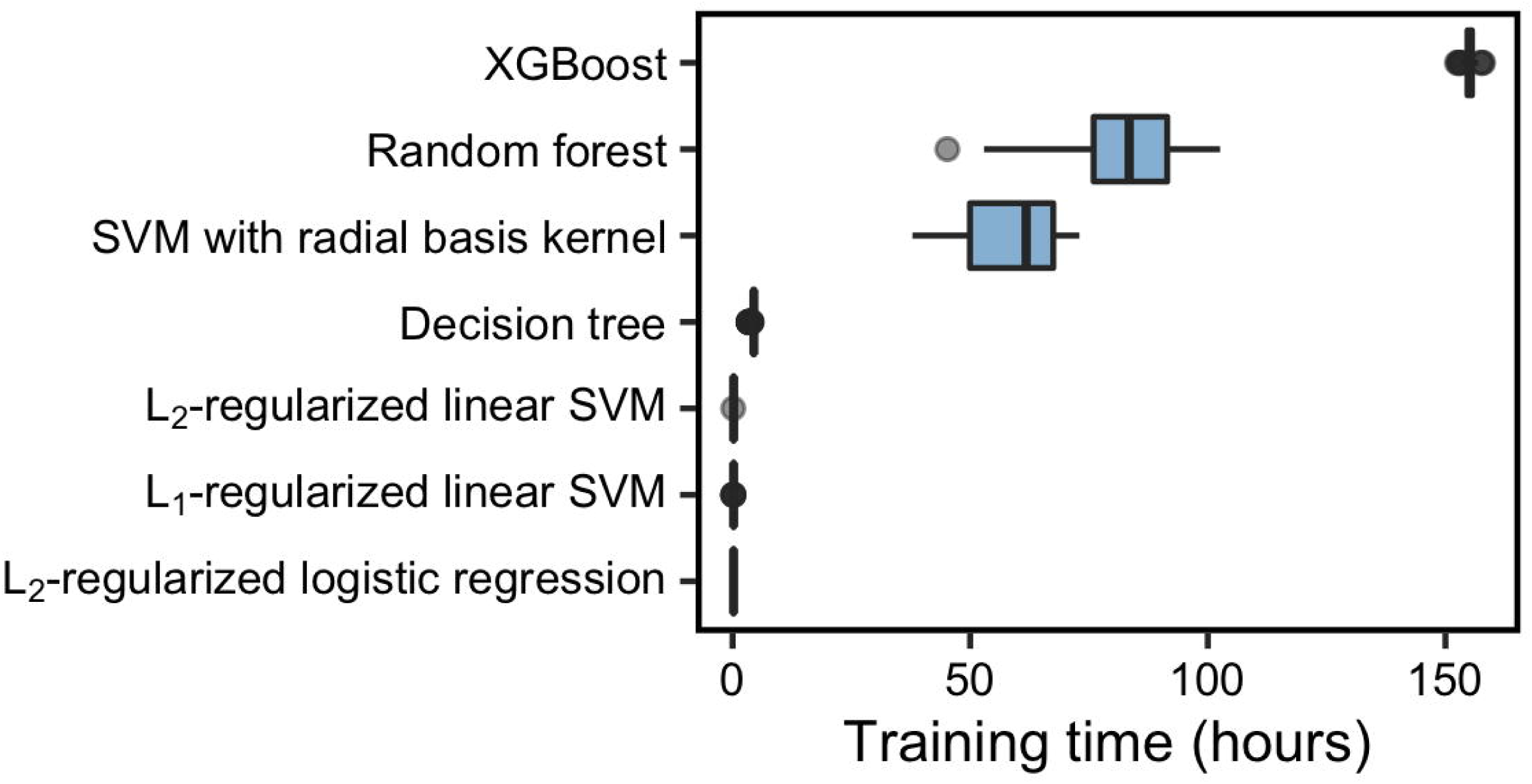
Training times of seven ML models. The median training time was the highest for XGBoost and shortest for L2-regularized logistic regression.

## Discussion

There is a growing awareness that many human diseases and environmental processes are not driven by a single organism but are the product of multiple bacterial populations. Traditional statistical approaches are useful for identifying those cases where a single organism is associated with a process. In contrast, ML methods offer the ability to incorporate the structure of the microbial communities as a whole and identify associations between community structure and disease state. If it is possible to classify communities reliably, then ML methods also offer the ability to identify those microbial populations within the communities that are responsible for the classification. However, the application of ML in microbiome studies is still in its infancy, and the field needs to develop a better understanding of different ML methods, their strengths and weaknesses, and how to implement them.

To address these needs, we developed an open-sourced framework for ML models. Using this pipeline, we benchmarked seven ML models and showed that the tradeoff between model complexity and performance may be less severe than originally hypothesized. In terms of predictive performance, the random forest model had the best AUROC compared to the other six models. However, the second-best model was L2-regularized logistic regression with a median AUROC difference of less than 0.015 compared to random forest. While our implementation of random forest took 83.2 hours to train, our L2-regularized logistic regression trained in 12 minutes. In terms of interpretability, random forest is a non-linear ML model, while L2-regularized logistic regression, a linear model, was more easily interpreted because we could use the feature weights. Comparing many different models showed us that the most complex model was not necessarily the best model for our ML task.

We established a pipeline that can be generalized to any modeling method that predicts a binary health outcome. We performed a random data-split to create a training set (80% of the data) and a held-out test set (20% of the data), which we used to evaluate predictive performance. We used the AUROC metric to evaluate predictive performance as it is a clinically relevant evaluation metric for our study. We repeated this data-split 100 times to measure the possible variation in predictive performance. During training, we tuned the model hyperparameters with a repeated five-fold cross-validation. Despite the high number of features microbiome datasets typically have, the models we built with this pipeline generalized to the held-out test sets.

We highlighted the importance of model interpretation to gain greater biological insights into microbiota-associated diseases. In this study, we showcased two different interpretation methods: ranking each OTU by (i) their absolute weights in the trained models and (ii) their impact on the predictive performance based on permutation importance. Previous studies have emphasized the difficulty of interpreting the feature coefficients in linear models (37) and the biases introduced by computing feature importance using built-in methods (e.g., gini drop) of tree-based models (38). Therefore, we encourage our audience to use both interpretation methods highlighted in this study as permutation importance is a model-agnostic tool that can be used to compared feature importance across different models. Human-associated microbial communities have complex correlation structures that create collinearity in the datasets. This can hinder our ability to reliably interpret models because the feature weights of correlated OTUs are influenced by one another (39). To capture all important features, once we identify highly ranked OTUs, we should review their relationships with other OTUs. These relationships will help us generate new hypotheses about the ecology of the disease and test them with follow-up experiments. When we used permutation importance, we partially accounted for collinearity by grouping correlated OTUs to determine their impact as a group. We grouped OTUs that had a perfect correlation with each other; however, we could reduce the correlation threshold to further investigate the relationships among correlated features. By our approach, we identified 432 OTUs out of 6,920 that had perfect correlations with at least one other OTU. The decision to establish correlation thresholds is left to researchers to implement for their own analyses. Regardless of the threshold, undestanding the correlation structures within the data is critical to avoid misinterpreting the models. Such structures are likely to be a particular problem with shotgun metagenomic datasets where collinearity will be more pronounced due to many genes being correlated with one another because they come from the same chromosome. Finally, true causal mechanisms (e.g., role of microbiome in colorectal cancer) cannot be explained solely by the highest performing machine learning model (40). To identify the true underlying microbial factors of a disease, it is crucial to follow up on any correlation analyses with further hypothesis testing and experimentation for biological validation.

In this study, we did not consider all possible modeling approaches. However, the principles highlighted throughout this study apply to other ML modeling tasks with microbiome data. For example, we did not evaluate multicategory classification methods to predict non-binary outcomes. We could have trained models to differentiate between people with healthy colons and those with adenomas or carcinomas (k=3 categories). We did not perform this analysis because the clinically relevant diagnosis grouping was between patients with healthy colons and those with SRNs. Furthermore, as the number of classes increases, more samples are required for each category to train an accurate model. We also did not use regression-based analyses to predict a non-categorical outcome. We have previously used such an approach to train random forest models to predict fecal short-chain fatty acid concentrations based on microbiome data (41). Our analysis was also limited to shallow learning methods and did not explore deep learning methods such as neural networks. Deep learning methods hold promise (12, 42, 43), but microbiome datasets often suffer from having many features and small sample sizes, which result in overfitting.

Our framework provides a reproducible structure for investigators wanting to train, evaluate, and interpret their own ML models to generate hypotheses regarding which OTUs might be biologically relevant. However, deploying microbiome-based models to make clinical diagnoses or predictions is a significantly more challenging and distinct undertaking (44). For example, we currently lack standardized methods to collect patient samples, generate sequence data, and report clinical data. We are also challenged by the practical constraints of OTU-based approaches. The de novo algorithms commonly in use are slow, require considerable memory, and result in different OTU assignments as new data are added (45). Finally, we also need independent validation cohorts to test the performance of a diagnostic model. To realize the potential for using ML approaches with microbiome data, it is necessary that we direct our efforts to overcome these challenges.

Our study highlights the need to make educated choices at every step of developing an ML model with microbiome data. We created an aspirational rubric that researchers can use to identify potential pitfalls when using ML in microbiome studies and ways to avoid them [Table S1]. We highlighted the trade-offs between model complexity and interpretability, the need for tuning hyperparameters, the utility of held-out test sets for evaluating predictive performance, and the importance of considering correlation structures in datasets for reliable interpretation. We showed the importance of interpretability for generating hypotheses to identify causal, biological relationships and for identifying inconsistencies in model setup. Furthermore, we underscored the importance of proper experimental design and methods to help us achieve the level of validity and accountability we want from models built for patient health.

## Materials and Methods

### Data collection and study population

The original stool samples described in our analysis were obtained from patients recruited by Great Lakes-New England Early Detection Research Network (5). Stool samples were provided by adults who were undergoing a scheduled screening or surveillance colonoscopy. Participants were recruited from Toronto (ON, Canada), Boston (MA, USA), Houston (TX, USA), and Ann Arbor (MI, USA). Patients’ colonic health was visually assessed by colonoscopy with bowel preparation and tissue histopathology of all resected lesions. We assigned patients into two classes: those with healthy colons and those with screen relevant neoplasias (SRNs). The healthy class included patients with healthy colons or non-advanced adenomas, whereas the SRN class included patients with advanced adenomas or carcinomas (46). Patients with an adenoma greater than 1 cm, more than three adenomas of any size, or an adenoma with villous histology were classified as having advanced adenomas (46). There were 172 patients with normal colonoscopies, 198 with adenomas, and 120 with carcinomas. Of the 198 adenomas, 109 were identified as advanced adenomas. Together 261 patients were classified as healthy and 229 patients were classified as having an SRN.

### 16S rRNA gene sequencing data

Stool samples provided by the patients were used for 16S rRNA gene sequencing to measure bacterial population abundances. The sequence data used in our analyses were originally generated by Baxter et al. (available through NCBI Sequence Read Archive [SRP062005], (5)). The OTU abundance table was generated by Sze et al (47), who processed the 16S rRNA sequences in mothur (v1.39.3) using the default quality filtering methods, identifying and removing chimeric sequences using VSEARCH, and assigning to OTUs at 97% similarity using the OptiClust algorithm (45, 48, 49); (https://github.com/SchlossLab/Sze_CRCMetaAnalysis_mBio_2018/blob/master/data/process/baxter/baxter.0.03.subsample.shared). These OTU abundances were the features we used to predict colorectal health of the patients. There were 6920 OTUs. OTU abundances were subsampled to the size of the smallest sample and normalized across samples such that the highest abundance of each OTU would be 1, and the lowest would be 0.

### Model training and evaluation

Models were trained using the caret package (v.6.0.81) in R (v.3.5.0). We modified the caret code to calculate decision values for models generated using L2-regularized SVM with linear kernel and L1-regularized SVM with linear kernel. The code for these changes on L2-regularized SVM with linear kernel and L1-regularized SVM with linear kernel models are available at https://github.com/SchlossLab/Topcuoglu_ML_XXX_2019/blob/master/data/caret_models/svmLinear3.R and at https://github.com/SchlossLab/Topcuoglu_ML_XXX_2019/blob/master/data/caret_models/svmLinear4.R, respectively.

For hyperparameter selection, we started with a granular grid search. Then we narrowed and fine-tuned the range of each hyperparameter. For L2-regularized logistic regression, L1 and L2- regularized SVM with linear and radial basis function kernels, we tuned the cost hyperparameter, which controls the regularization strength, where smaller values specify stronger regularization. For SVM with radial basis function kernel, we also tuned the sigma hyperparameter, which determines the reach of a single training instance where, for a high value of sigma, the SVM decision boundary will be dependent on the points that are closest to the decision boundary. For the decision tree model, we tuned the depth of the tree where the deeper the tree, the more splits it has. For random forest, we tuned the number of features to consider when looking for the best tree split. For XGBoost, we tuned the learning rate and the fraction of samples used for fitting the individual base learners. Performing a grid search for hyperparameter selection might not be feasible for when there are more than two hyperparameters to tune for. In such cases, it is more efficient to use random search or recently developed tools such as Hyperband to identify good hyperparameter configurations (50).

The computational burden during model training due to model complexity was reduced by parallelizing segments of the ML pipeline. We parallelized the training of each data-split. This allowed the 100 data-splits to be processed through the ML pipeline simultaneously at the same time for each model. It is possible to further parallelize the cross-validation step for each hyperparameter setting which we have not performed in this study.

### Permutation importance workflow

We calculated a Spearman’s rank-order correlation matrix and defined correlated OTUs as having perfect correlation (correlation coefficient = 1 and p < 0.01). OTUs without a perfect correlation to each other were permuted individually, whereas correlated ones were grouped together and permuted at the same time.

### Statistical analysis workflow

Data summaries, statistical analysis, and data visualizations were performed using R (v.3.5.0) with the tidyverse package (v.1.2.1). We compared the performance of the models pairwise by calculating the difference between AUROC values from the same data-split (for 100 data-splits). We determined if the models were significantly different by calculating the empirical p-value (2 x min(% of AUROC differences ≥ 0, % of AUROC differences < 0) for the double tail event (e.g., Figure S4).

### Code availability

The code for all sequence curation and analysis steps including an Rmarkdown version of this manuscript is available at https://github.com/SchlossLab/Topcuoglu_ML_XXX_2019/.

## Acknowledgements

We thank all the study participants of Great Lakes-New England Early Detection Research Network. We would like to thank the members of the Schloss lab for their valuable feedback. Salary support for MR came from NIH grant 1R01CA215574. Salary support for PDS came from NIH grants P30DK034933 and 1R01CA215574.

---

**Figure S1. NMDS ordination of Bray-Curtis distances**. NMDS ordination relating the community structures of the fecal microbiota from 490 patients (261 patients with normal colonoscopies and 229 patients who have screen relevant neoplasias; SRNs).

**Figure S2. Hyperparameter setting performances for linear models**. (A) L2-regularized logistic regression, (B) L1-regularized SVM with linear kernel, and (C) L2-regularized SVM with linear kernel mean cross-validation AUROC values when different hyperparameters were used in training the model. The stars represent the highest performing hyperparameter setting for each model.

**Figure S3. Hyperparameter setting performances for non-linear models**. (A) Decision tree, (B) random forest, (C) SVM with radial basis kernel, and (D) XGBoost mean cross-validation AUROC values when different hyperparameters were used in training the model. The stars represent the highest performing hyperparameter setting for the models.

**Figure S4. Histogram of AUROC differences between L2-regularized logistic regression and random forest for each of the hundred data-splits**. In 75% of data-splits, the AUROC of random forest was greater than that of L2-regularized logistic regression. The p-value was manually calculated using the sampling distribution of the test statistic under the null hypothesis. We tested how often random forest performed more accurately than L2-regularized logistic regression. The null hypothesis is that the distribution of the difference between the AUROC values of random forest and L2 logistic regression is symmetric about 0, therefore the p-value was calculated for a double-tail event.

**Figure S5. Classification performance of ML models across cross-validation when trained on a subset of the dataset**. (A) L2-regularized logistic regression and (B) random forest models were trained using the original study design with 490 subjects and subsets of it with 15, 30, 60, 120, and 245 subjects. The range among the cross-validation AUROC values within both models at smaller sample sizes were much larger than when the full collection of samples was used to train and validate the models but included the ranges observed with the more complete datasets.

**Figure S6. Permutation importance analysis**. Permutation importance analysis measures the decrease in the predictive performance of the model after we permute (A) a feature’s or (B) a group of correlated features’ values, which breaks the relationship between the feature and the diagnosis.

**Figure S7. Interpretation of the linear ML models with permutation importance**. (A) L1- regularized SVM with linear kernel, (B) L2-regularized SVM with linear kernel, and (C) L2- regularized logistic regression were interpreted using permutation importance using held-out test set.

**Figure S8. Relative abundances of the 20 most important OTUs in L2-regularized logistic regression and random forest models**. The most important 20 OTUs were chosen for (A) Random forest and (B) L2-regularized logistic regression models by permutation importance and ranking feature coefficients, respectively. The relative abundances of these OTUs were compared based on the diagnosis of the patients. The minimal differences betweeen relative abundances for these OTUs show that it is not possible to differentiate disease vs healthy states by focusing on individual taxa.

